# Lost in the WASH. The functional human WASH complex 1 gene is on chromosome 20

**DOI:** 10.1101/2023.06.14.544951

**Authors:** Daniel Cerdán-Vélez, Michael L. Tress

## Abstract

The WASH1 gene produces a protein that forms part of the developmentally important WASH complex. The WASH complex activates the Arp2/3 complex to initiate branched actin networks at the surface of endosomes. As a curiosity, the human reference gene set includes nine WASH1 genes. How many of these are pseudogenes and how many are *bona fide* coding genes is not clear.

Eight of the nine WASH1 genes reside in rearrangement and duplication-prone subtelomeric regions. Many of these subtelomeric regions had gaps in the GRCh38 human genome assembly, but the recently published T2T-CHM13 assembly from the Telomere to Telomere (T2T) Consortium has filled in the gaps. As a result, the T2T Consortium has added four new WASH1 paralogues in previously unannotated subtelomeric regions.

Here we show that one of these four novel WASH1 genes, *LOC124908094*, is the gene most likely to produce the functional WASH1 protein. We also demonstrate that the other twelve WASH1 genes derived from a single *WASH8P* pseudogene on chromosome 12. These 12 genes include WASHC1, the gene currently annotated as the functional WASH1 gene.

We propose *LOC124908094* should be annotated as a coding gene and all functional information relating to the *WASHC1* gene on chromosome 9 should be transferred to *LOC124908094*. The remaining WASH1 genes, including *WASHC1*, should be annotated as pseudogenes. This work confirms that the T2T assembly has added at least one functionally relevant coding gene to the human reference set. It remains to be seen whether other important coding genes are missing from the GRCh38 reference assembly.

## Introduction

The Arp2/3 complex has a crucial role in the generation of branched actin filaments that are important in a range of processes including endocytosis, intracellular trafficking and membrane remodelling. The WASH (Wiskott-Aldrich syndrome protein and SCAR Homologue) complex is one of a number of complexes that activate the Arp2/3 complex at different subcellular locations [Alekhine et al, 2017]. The WASH complex plays a key role in endosome sorting by recruiting and activating the Arp2/3 complex to induce actin polymerization [Derivery et al, 2009, Gomez and Billadeau, 2009]. The WASH complex has been shown to be essential in both Drosophila [Linardopoulou et al, 2007] and mice [Xia et al, 2014], and defects in the components of the WASH complex have been found to lead to inherited developmental and neurological disorders [Valdmanis et al, 2007, Courtland et al, 2021].

The WASH complex is made up of proteins from five genes, *WASHC1* (previously *WASH1*), *WASHC2A* (*FAM21A*), *WASHC3* (*CCDC53*), *WASHC4* (*SWIP*) and *WASHC5* (*KIAA0196*). The exact composition of the complex is complicated by the fact that there are two highly similar FAM21 genes with peptide support, *WASHC2A* and *WASHC2C*, and because great apes, and in particular humans, have multiple WASH1 genes.

Most species have a single *WASHC1* gene, but in great apes *WASH1* genes are found in rearrangement-prone subtelomeric regions near the ends of chromosomes [Linardopoulou et al, 2007]. Gene duplications from interchromosomal duplication are common in subtelomeric regions, and gorilla, chimpanzee and human all have multiple copies of the WASH1 gene [Linardopoulou et al, 2007]. In the human genome, there are nine copies of WASH1 annotated in the GRCh38 human reference set: two pseudogenes on the p arm of chromosome 1, and seven copies annotated on chromosomes 2, 9, 12, 15, 16 and 19, and the pseudoautosomal regions of X and Y. Eight are found in subtelomeric regions, while the ninth, *WASH2P*, is present in ancestral telomere-telomere fusion site on chromosome 2 [IJdo et al, 1991].

Many of the nine WASH1 copies currently annotated in the human reference gene set are either not full length, or have premature stop codons or frameshifts that will likely render them inactive [Linardopoulou et al, 2007]. Just one copy is full length and intact, the copy on chromosome 9, and this gene is thought to form part of the WASH complex. It is annotated as *WASHC1* (WASH complex subunit 1). The other eight genes are mostly annotated as pseudogenes, and have gene identifiers that mark them out as pseudogenes, from *WASH2P* (on chromosome 2) to *WASH9P* (chromosome 1).

There is some disagreement between the three main reference databases as to whether genes other than *WASHC1* can code for proteins. RefSeq [Sayers et al, 2023a] annotates *WASHC1* alone as coding, but Ensembl/GENCODE [Frankish et al, 2023, Martin et al, 2023] predicts that both *WASHC1* and *WASH6P* (chromosomes X/Y) are coding. UniProtKB [The UniProtKB Consortium, 2023] lists proteins for *WASH2P, WASH6P, WASH3P* (chromosome 15) and *WASH4P* (chromosome 16), as well as for *WASHC1. WASH4P* and *WASH6P* lost their 5’ ends when duplicated, so have novel 5’ ends. Meanwhile, *WASH2P* contains two premature stop codons, and *WASH3P* has a 4-base frame-shifting deletion and a premature stop codon.

Part of the reason for the discrepancies between the reference sets is that a lot of the peptide evidence from proteomics experiments maps to other predicted WASH1 proteins and not to *WASHC1*. This suggests the existence of distinct WASH1 isoforms produced from different genes. The WASH1 cDNA (BC048328.1) produced by the Mammalian Gene Collection Program [Strausberg et al, 2002] is also quite distinct from the cDNA expected from *WASHC1*. Given the possible confusion among the various WASH1 paralogues, it is notable that experiments investigating the function of the WASH complex and WASH1 have tended not to use human *WASHC1*. For example, the initial experiments that discovered the function of the WASH complex used murine WASH1 [Derivery et al, 2009], or the product of the BC048328.1 cDNA [Linardopoulou et al, 2007, Gomez and Billadeau, 2009].

The ancestral WASH1 gene in apes is predicted to be the copy located on the p arm of chromosome 12. Probes used in distinct primate species detected WASH1 genes here in macaque, orangutan, gorilla and chimpanzee as well as human [Linardopoulou et al, 2007]. In addition, multiple WASH1 paralogues were detected in subtelomeric regions in the gorilla, chimpanzee and human genomes [Linardopoulou et al, 2007]. Along with the ancestral gene on chromosome 12 (*WASH8P* in the human genome), a second WASH1 gene on the p arm of chromosome 20 was detected in all three species [Linardopoulou et al, 2007]. This gene was not present in the GRCh38 assembly of the human genome.

With the publication of the T2T-CM13 genome assembly by the Telomere-to-Telomere consortium [Nurk et al, 2022], which has filled in the remaining 8% of the human genome and added 1,956 gene predictions to the GRCh38 reference, there are now at least four new predicted WASH1 paralogues. These are *LOC124906207* on chromosome 3, *LOC124906931* on chromosome 11, and *LOC124908094* and *LOC124908102* at opposite ends of chromosome 20. The two novel genes on chromosome 20 could produce full length proteins if they were translated. Here we show that one of these two novel full length WASH1 genes is likely to produce the WASH1 isoform that forms part of the WASH complex.

## Methods

### Novel sequences

Predicted protein sequences for *WASHC1, WASH2P, WASH3P, WASH4P* and *WASH6P* were obtained from UniProtKB. The predicted protein sequences of XP_047302880.1 and XP_047302896.1 from *LOC124908094* and *LOC124908102* were downloaded from RefSeq.

The DNA sequences of transcripts XR_007073267.1, XM_047446924.1, XM_047446940.1 from *LOC124906207, LOC124908094* and *LOC124908102* were downloaded from RefSeq. Transcript sequences with the equivalent exon combinations for *WASHC1, WASH2P, WASH3P, WASH7P, WASH8P* and *WASH9P*, and for chimpanzee and bonobo WASH1 genes on chromosomes 12 and 20 were downloaded from the Ensembl or RefSeq databases.

We downloaded the protein sequence generated from clone BC048328.1 from GenBank [Sayers et al, 2023b].

### Peptide mapping

We downloaded the full list of peptides detected for WASH1 genes that were recorded in the current version (2023-01) of the PeptideAtlas database [Kusenbach et el, 2014]. PeptideAtlas records peptides that map to *WASHC1* and to predicted pseudogenes *WASH2P, WASH3P, WASH4P* and *WASH6P* because UniProtKB predicts that these genes produce proteins. However, PeptideAtlas also detects multiple peptides that do not map to any of these sequences, so appears to have searched against other predicted WASH1 proteins in the past.

PeptideAtlas carries out a reanalysis of spectra from multiple large- and small-scale proteomics experiments using state-of-the-art search engines before post-processing using the in-house trans-proteomic pipeline [Deutsch et al, 2023]. Each peptide may be detected multiple times over the distinct experiments. The number of distinct experiments in which a peptide is detected is recorded as “observations” in the PeptideAtlas record for each peptide. We used these observations as a rough measure of the expression level of each peptide.

To be sure that we were working with the best quality peptides from PeptideAtlas, we required that all peptides had at least two observations (we removed all singletons), that the peptides were fully tryptic, and that the peptides has no more than two missed lysine or arginine cleavages.

We remapped the peptides recorded in PeptideAtlas to the proteins generated by the five genes annotated as proteins in UniProtKB, and to the predicted sequence of XP_047302880.1 from gene *LOC124908094*. This gene is located on chromosome 20 at the terminus of the p arm of chromosome 20. The predicted protein XP_047302880.1 is referred to throughout the paper as WASH1-20p13.

We had to remap the peptides to the WASH1 sequences from UniProtKB because many peptides that do not correspond to the *WASHC1* protein sequence were erroneously annotated as mapping to the *WASHC1* protein.

### Phylogenetic analysis

For the WASH1 phylogenetic tree we included the full length WASH1 genes annotated in human, chimpanzee and bonobo. These included six of the nine WASH1 genes annotated in the GRCh38 assembly of the human genome (*WASH4P, WASH5P* and *WASH6P* were left out of the analysis because they are missing exons) and three of four WASH1 genes added in the T2T-CM13 assembly (the gene on chromosome 11 was left out because no transcript was annotated with all the coding exons). Both chimpanzee and bonobo are annotated with two WASH1 genes, in both species the transcripts are full length and located on chromosomes 12 and 20. The phylogenetic tree of the WASH1 genes was generated with Phyml 3.0 [Guindon et al, 2010] using the TN93 +I substitution model according to SMS [Lefort et al, 2017] and running 1000 non-parametric bootstrap replicas. Chimpanzee and bonobo were forced to root the tree in the analysis.

## Results

### Annotated WASH1 genes have a similar gene neighbourhood

All but one of the 9 previously known WASH1 genes, those annotated in the gene set from the GRCh38 build, are located at the subtelomeric regions at the terminal end of chromosomes [Figure 1A]. Subtelomeric regions are prone to rearrangement and interchromosomal duplications [Linardopoulou et al, 2005]. As a result, gene content in subtelomeric regions has changed substantially since humans and chimpanzees split, and interchromosomal duplications have led to the generation of multiple WASH1 paralogues.

**Figure 1.**
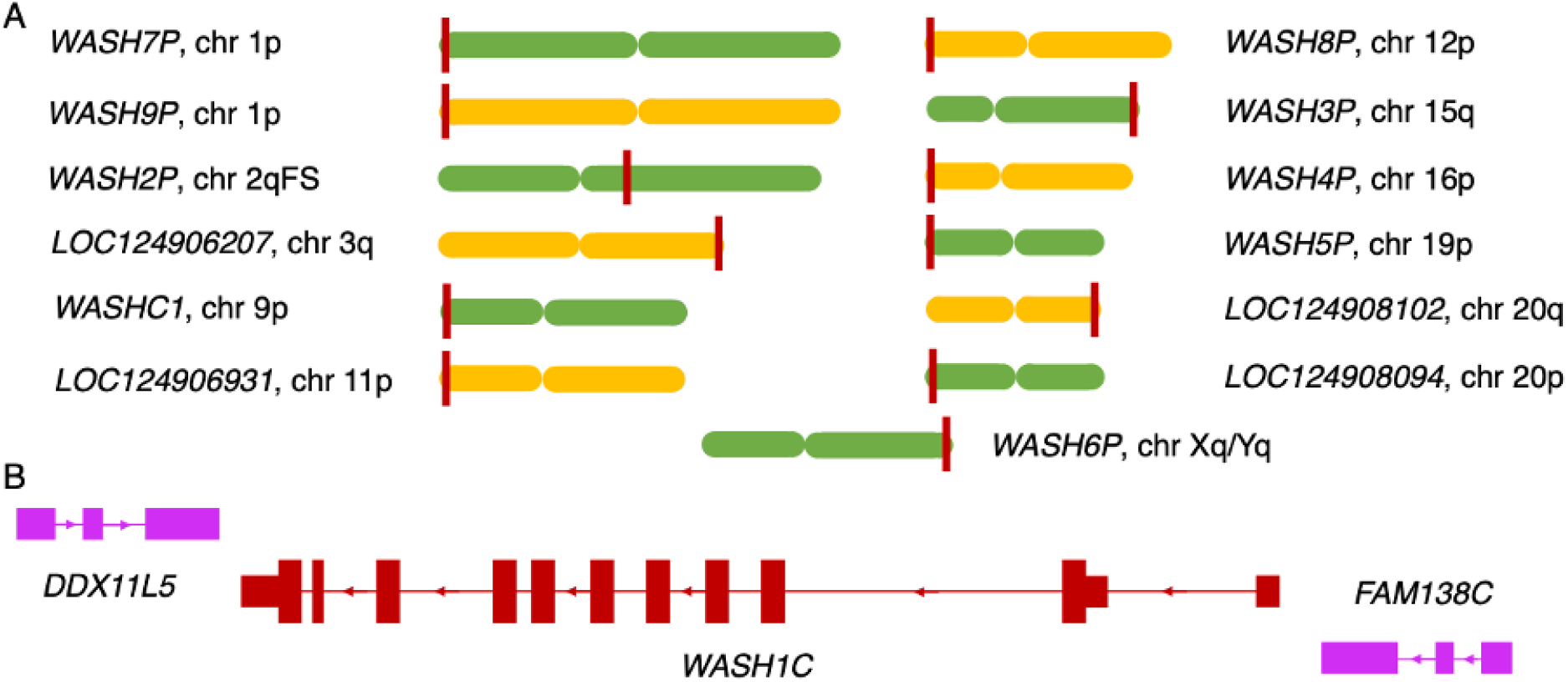
Annotated human WASH1 genes and their gene neighbourhood. A. The 13 WASH1 genes in the human genome and their approximate position in the chromosome. Chromosomes are represented by a single chromatid, alternate green and yellow and are not drawn to scale. The approximate position of the labelled WASH1 gene is indicated with a dark red vertical line. WASH1 genes are labelled by whether they are found on the p arm (p), the q arm (q) or the ancestral fusion site (FS). The p arm of chromosome 1 has two WASH1 genes on different strands. Genes annotated for the first time in the T2T assembly have names that start with “LOC”. B. The gene neighbours of gene WASHC1 on chromosome 9. Exons and introns not to scale. Exons shown as blocks. Coding exons are wider than non-coding exons. Coding genes are dark red. Non-coding genes/pseudogenes pink. Direction of the strand is shown by the arrows in the intronic regions.

The nature of interchromosomal duplication means that the multiple WASH1 genes were generated as part of the duplication of large blocks of subtelomeric regions, and these included the neighbouring genes. That means that most WASH1 genes have similar gene neighbours. Even those genes where only part of the WASH1 gene was duplicated (*WASH4P, WASH5P, WASH6P*) have retained their gene neighbours on one side at least.

Most WASH1 genes are preceded upstream on the same strand by a gene from the FAM138 family [Figure 1B]. FAM138 genes are non-coding in the human gene set, but are annotated as coding in some primate species. Just upstream of the WASH1 genes, and on the opposite strand, is a DDX11-like gene, a probable pseudogene made up of a short fragment of a dead-box helicase gene. Curiously, the association between WASH1 and DDX11-like helicases has ancient roots. Zebrafish has a single *wash1* gene, and the coding gene on the opposite strand downstream of *wash1* is the full length *ddx11* gene. When WASH1 migrated to the subtelomeric regions in primates it only took part of the *DDX11* gene with it.

The gene neighbourhood is the same for all WASH1 genes in the GRCh38 reference genome except for *WASH5P*, which lies adjacent to block of missing sequence on chromosome 19, and *WASH4P* and *WASH6P*, which were both duplicated without their 5’ exons and as a result lost the FAM138 family gene too. Instead *WASH4P* and *WASH6P* have downstream WASIR genes.

The four previously unannotated WASH1 genes uncovered in the T2T-CHM13 human genome assembly are on chromosomes 3, 11, and 20 [Figure 1A]. Genes *LOC124906207, LOC124906931* and *LOC124908102* have the same gene neighbourhood as the other full length genes in GRCh38, an upstream FAM138 family gene and a downstream DDX11-like gene, but *LOC124908094*, on the p arm of chromosome 20, is missing the FAM138 family gene.

There are multiple duplications of WASH1 genes in all great apes [Linardopoulou et al, 2007]. The reference sets of gorilla, bonobo and chimpanzee have two copies of WASH1 genes each. In gorilla, WASH1 genes are annotated on chromosomes 3 and 6, while in chimpanzee and bonobo they are annotated on chromosomes 12 and 20. In chimpanzee, the chromosome 12 WASH1 gene neighbourhood is bounded by both a DDX11-like pseudogene and a FAM138 family gene, just like the equivalent human WASH1 gene on chromosome 12. The WASH1 copy on chromosome 20 in chimpanzee is adjacent just to the DDX11-like pseudogene. As with the WASH1 gene on the p arm of chromosome 20 in human, there is no upstream FAM138 family gene.

### Peptide evidence supports a single WASH1 gene

In order to determine which of the WASH1 genes might be coding, we mapped all peptides recorded in PeptideAtlas to the five WASH1 isoforms annotated in UniProtKB and to the predicted sequence of the full length WASH1 protein identified on the p arm of chromosome 20, XP_047302880.1 (referred to as WASH1-20p13 in the remainder of the text).

In total, 52 peptides with a total of 23,812 observations in PeptideAtlas map to at least one of the six proteins. The peptides, coloured by the number of observations are shown mapped onto the sequences of the six predicted proteins in Figure 2. None of the proteins captures all 52 of the PeptideAtlas peptides, suggesting that more than one of the genes might be coding. The only gene annotated as coding in the three main human reference databases, *WASHC1*, performs particularly poorly. Only 26 of the 52 PeptideAtlas peptides map to the protein produced by *WASHC1*. This protein is supposed to have a functional role as a component of the nucleation-promoting endosome-based WASH core complex yet just half of the reliable tryptic peptides from PeptideAtlas map to the *WASHC1* gene product.

**Figure 2.**
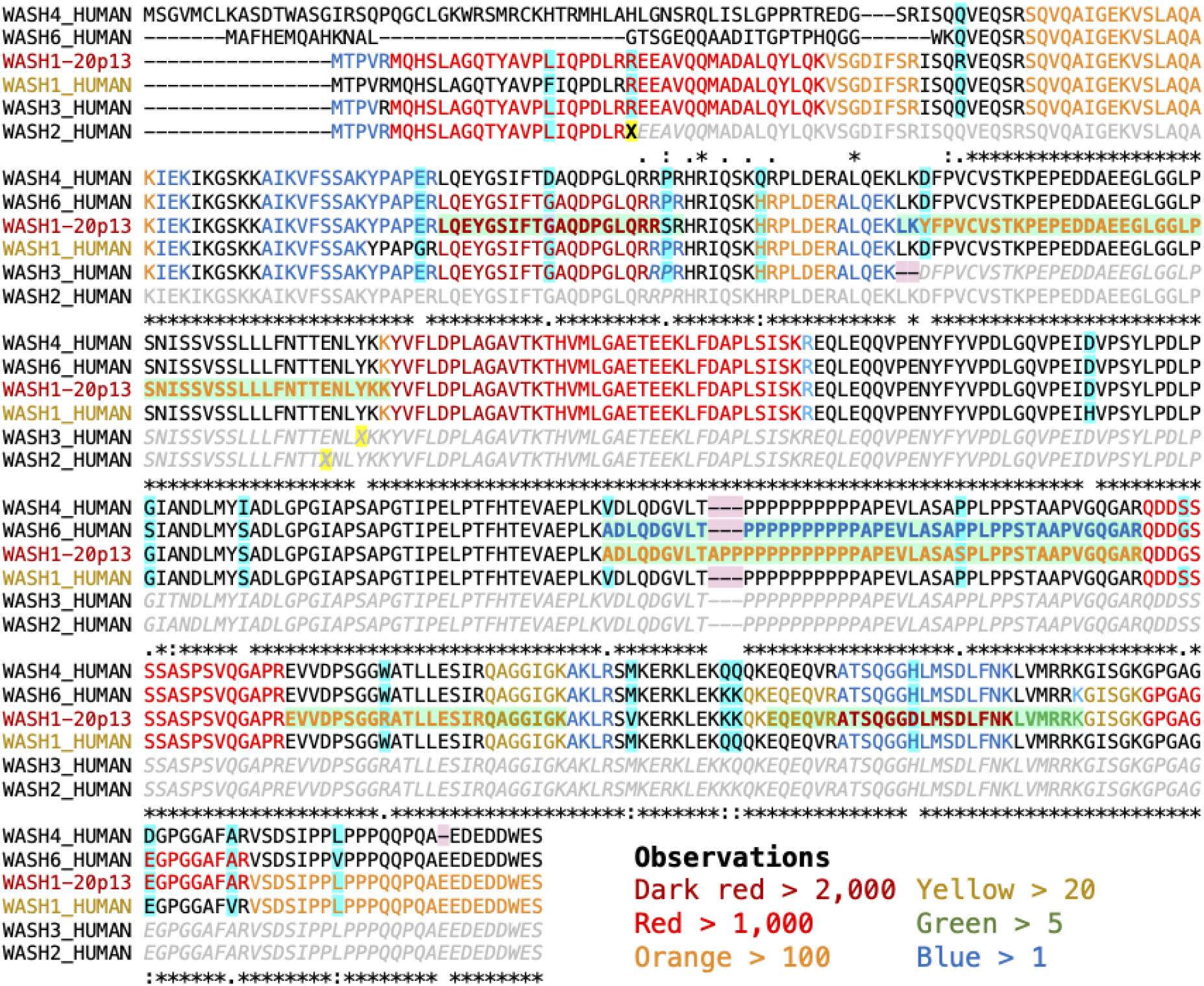
Mapping peptides to WASH1-20p13 and WASH1 sequences in UniProtKB. Peptides are mapped to alignments between the five UniProtKB annotated WASH1 sequences and the newly annotated WASH1-20p13 protein. Residues that peptides map to are colour coded by the number of observations detected for that protein. Dark red, red and orange font indicate the most observed peptides. Residues with a yellow background indicate the position of stop codons and frameshifts in *WASH2P* and *WASH3P*. Grey sequence indicates the parts of the predicted sequence of *WASH2P* and *WASH3P* that cannot be translated because of these stop codons and frameshifts. Peptides with a light green background and text in bold show those peptides that map uniquely to one of the six sequences. Amino acids columns with a blue background indicate the position of single amino acid differences between the predicted proteins. Deletions relative to ancestral sequences are marked with a pink background.

Ensembl/GENCODE annotates *WASH6P* and *WASHC1* as coding. However, *WASH6P* and *WASHC1* between them capture only 34 PeptideAtlas peptides. This still leaves more than a third of PeptideAtlas WASH1 peptides unaccounted for. Extending the mapping to the five predicted proteins in UniProtKB (adding predicted proteins from *WASH2P, WASH3P* and *WASH4P* to *WASHC1* and *WASH6P*) would cover most of the 52 WASH1 peptides in PeptideAtlas, if the predicted sequences of *WASH2P* and *WASH3P* in UniProtKB were correct. Unfortunately, these predictions are false. *WASH3P* has a frameshift that would eliminate the final two thirds of the protein, and *WASH2P* has two premature stop codons, the first of which would leave a rump of just 25 amino acid residues. If translated, the truncated proteins produced by these two genes would capture just 21 and 2 PeptideAtlas peptides, but both of these genes are highly likely to be pseudogenes.

By way of contrast, the *LOC124908094* predicted protein, WASH1-20p13, captures 47 of the 52 PeptideAtlas peptides. Twelve of the peptides map to WASH1-20p13 and no other protein, including one of the three most highly observed peptides, ATSQGGDLMSDLFNK, with more than 2,000 observations in PeptideAtlas. If we assume that pseudogenes *WASH2P* and *WASH3P* do not produce their truncated proteins, another five peptides would map just to the WASH1-20p13 isoform, including the peptide MQHSLAGQTYAVPLIQPDLR that has 1,300 observations. Together the 17 peptides that map to WASH1-20p13 alone strongly suggest that *LOC124908094* is the gene that produces the WASH complex protein.

We also mapped peptides to the predicted isoform of the other full length WASH1 gene identified in the T2T assembly on chromosome 20, XP_047302896.1. However, XP_047302896.1 had little support; just 17 of the 52 PeptideAtlas peptides mapped to this protein.

With the peptides weighted by the number of observations from the peptides that map to them, the importance of WASH1-20p13 becomes even more apparent. A total of 22,564 of the 23,813 observations from all the PeptideAtlas peptide map to WASH1-20p13, with the next highest total (15,592 observations) for the *WASHC1* protein [**Figure 3A**].

**Figure 3.**
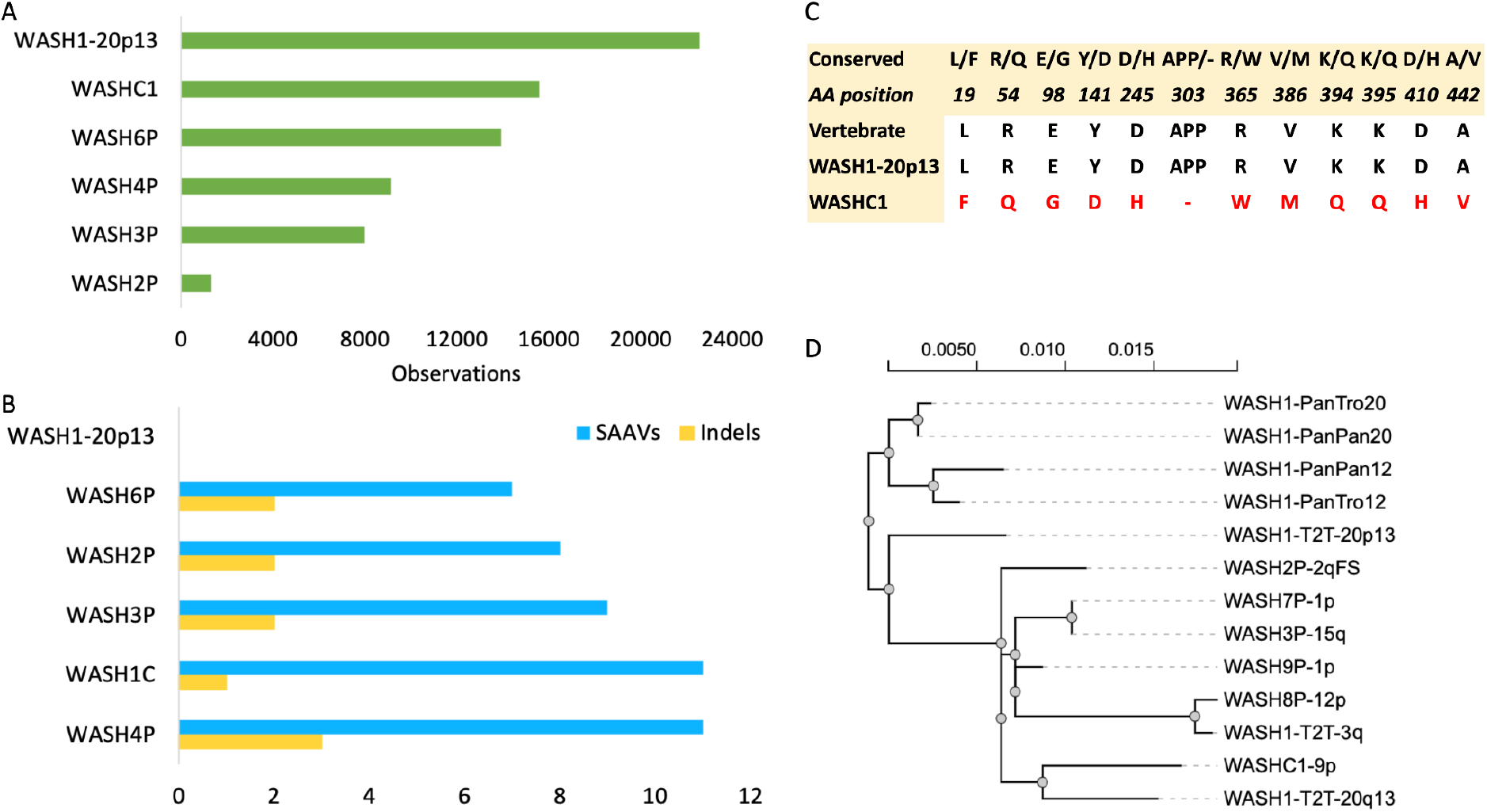
PSM observations, variants and phylogenetic analysis. A. The sum of the observations from all PSM that map to each of the five WASH1 isoforms annotated in UniProtKB and the protein from the newly annotated WASH1 gene on chromosome 20, WASH1-20p13. B. The count of non-conserved single amino acids and deleted regions that differ from the conserved amino acids across primates, mammals and tetrapods in the same six WASH1 isoforms. WASH1-20p13 is left blank because it has no changes in these highly conserved regions. C. The amino acid differences between the protein currently annotated as the functional WASH1 protein (from the *WASHC1* gene) and WASH1-20p13 in regions conserved across vertebrates. The *WASHC1* protein is different in all 12 conserved positions. D. The results of a phylogenetic analysis that includes annotated WASH1 genes for chimpanzee, bonobo and the nine annotated WASH1 genes that maintain the ancestral exons (*WASH4P, WASH5P, WASH6P* and *LOC124906931* are not annotated with all exons). Genes newly annotated in T2T assembly are labelled as WASH1-T2T plus their chromosome position. Bonobo (PanPan) and chimpanzee (PanTro) genes and known human genes are also labelled by the chromosome in which they are annotated.

### G343S appears to be a common variant in WASH1-20p13

Five of the 52 WASH1 peptides in PeptideAtlas do not map to WASH1-20p13. There are three possible explanations for these unmapped peptide spectrum matches (PSMs). Either two (or more) WASH1 genes are translated, the PSMs are false positive matches, or the WASH1-20p13 has one or more common amino acid variants. None of the other WASH1 proteins captures all five of these peptides. Four of the five peptides have four or fewer observations, but one peptide, QDDSSSSASPSVQGAPR, was detected 1,239 times in experiments across PeptideAtlas. A similar peptide, QDDGSSSASPSVQGAPR, which maps to WASH1-20p13, was observed 1,999 times.

It is curious that peptide QDDSSSSASPSVQGAPR has so many observations when none of the other four non-mapping peptides have more than 4. One possibility is that there is a common SNP in WASH1-20p13 that changes glycine residue 343 into a serine. It is of note that the only difference between WASH1-20p13 and the protein produced by the BC048328.1 cDNA clone from the Mammalian Gene Collection Programme is the same glycine to serine substitution.

If the G343S variant is a feature of the *LOC124908094* gene, 49 of the 52 PeptideAtlas peptides and 23,804 of the 23,813 observations would map to WASH1-20p13. The remaining 3 peptides, with a total of 9 observations, would all map to *WASH6P*.

### Cross-species conservation supports the role of WASH1-20p13 in the WASH complex

Alignments between the five WASH1 isoforms annotated in UniProtKB and WASH1-20p13 reveal 21 single amino acid differences and three deletions, as well as the distinct novel 5’ coding exons in *WASH4P* and *WASH6P* [Figure 2]. Multiple alignments of WASH1 proteins from vertebrate species show that 17 of these single residue differences are conserved across tetrapod WASH1 proteins. In all 17 single amino acid differences conserved across tetrapods, the WASH1-20p13 isoform has the conserved amino acid. None of the deletions are found outside of the human WASH1 paralogues, and it is also the only protein with none of the human-specific deletions [Figure 3B]. All the other five predicted isoforms have at least one indel that is different from the conserved vertebrate WASH1 sequence. The isoforms from *WASHC1* and *WASH4P* have eleven human-specific substitutions of amino acids that are conserved in mammals, birds and amphibians.

Most of the amino acid substitutions in annotated UniProtKB WASH1 proteins are not conservative. For example, just one of the 11 single amino acid differences between the *WASHC1* protein and WASH1-20p13 that fall in conserved regions might be considered conservative (the valine for alanine swap in residue 439 of the *WASHC1* protein). The full list of differences between the *WASHC1* protein and WASH1-20p13 can be seen in Figure 3C.

Several amino acid substitutions in *WASHC1* are considered radical. One example is the tyrosine to aspartate swap at residue 141. Tyrosine residue Y141 is phosphorylated by the Src kinase LCK and mutating this residue to phenylalanine has been shown to interfere with WASH-mediated NK cell cytotoxicity [Huang et al, 2016]. However, aspartate is a phosphomimetic, so a Y141D mutation may well lead to a permanent intensification of NK cell cytotoxicity. All twelve predicted WASH1 proteins other than WASH1-20p13 have this Y141D substitution.

### Is *LOC124908094* the only WASH1 coding gene?

There are 13 human WASH1 genes annotated in the T2T-CM13 assembly, which means that as well as the likely ancestral genes on chromosome 12 and 20 (*WASH8P* and *LOC124908094*), there are 11 novel WASH1 paralogues. Several are clearly pseudogenes. *WASH8P*, the ancestral copy of WASH1 on chromosome 12, has undergone many changes. *WASH8P* is no longer functional because it would generate a transcript with a premature stop codon. The same stop codon is present in four other human WASH1 pseudogenes, *WASH3P, WASH9P, WASH7P*, and *LOC124906207*, the copy of WASH1 annotated by the T2T consortium on chromosome 3.

Like humans, bonobo and chimpanzee have copies of WASH1 at the terminal end of the p arm of chromosome 12 (like human *WASH8P*) and at the terminal end of the p arm of chromosome 20 (like *LOC124908094*). Alignments between the annotated WASH1 isoforms in chimpanzee and bonobo reveal they have few amino acid residue differences (the isoform produced by WASH1 on chromosome 12 in bonobo has four substitutions relative to the other three isoforms; none of the other proteins have more than one), suggesting that in bonobo and chimpanzee at least, both ancestral paralogues might be functional coding genes.

The 3-residue deletion of amino acids “APP” is possibly the most interesting difference between WASH1-20p13 and the other potential WASH1 isoforms [Figure 2]. The three amino acids missing from the start of the proline-rich region in all five annotated WASH1 isoforms in UniProtKB are not lost in any other species, though lemurs have lost three other prolines from the proline-rich region. The amino acids AAP are conserved in all four WASH1 isoforms in chimpanzee and bonobo. In human, while WASH1-20p13 from *LOC124908094* maintains the three amino acids at the start of the proline-rich region, the predicted protein from WASH8P, the other ancestral copy of WASH1 protein, would not.

If translated, 10 of the 11 copies of WASH1 that duplicated from the ancestral WASH1 genes would have produced proteins with the three amino acid deletion in the proline-rich region.

The 11th, *WASH5P*, is highly truncated. This deletion and the multiple amino acid substitutions that are not present in any non-human WASH1 gene suggest that *WASHC1, WASH2P, WASH3P, WASH4P, WASH6P, WASH7P, WASH9P, LOC124908102, LOC124906931* and *LOC124906207* all arose in the human lineage.

There are clues that all duplications derived from *WASH8P* rather than *LOC124908094*. Nine of the 11 novel paralogues (including the truncated *WASH5P* gene) have an upstream FAM138 family pseudogene. FAM138 pseudogenes are found upstream of the chromosome 12 WASH1 genes in both human and chimpanzee, but not upstream of the chromosome 20 WASH1 genes. The only paralogues without an upstream FAM138-like pseudogene are *WASH4P* and *WASH6P*, but this is not surprising since both also lost their 5’ coding exons when they duplicated.

The phylogenetic analysis of the ancestral WASH1 genes on chromosomes 12 and 20 (two genes each from chimpanzee and bonobo along with human *WASH8P* and *LOC124908094*) and the seven human WASH1 duplications that are full length provides the final confirmation that these novel paralogues derived in the human lineage and from *WASH8P* rather than *LOC124908094* [Figure 3D].

Although the accumulated evidence clearly suggests that all 11 novel WASH1 paralogues derived from *WASH8P*, not all duplicated directly from the *WASH8P* pseudogene. *LOC124906207*, the copy of WASH1 annotated on chromosome 3 by the T2T Consortium, is a recent duplication of *WASH8P*, but *WASH3P, WASH7P* and WASH9P appear to have a single common ancestor that itself duplicated from *WASH8P. WASH4P* and *WASH6P* are also likely to have a common ancestor that duplicated from *WASH8P* given the loss of their 5’ exons and their similar gene neighbourhoods.

## Conclusions

The new T2T assembly of the human genome has added 4 new WASH1 genes to the 9 that were already known. Many of these WASH1 paralogues appear to be pseudogenes. Until now, *WASHC1* on chromosome 9 was believed to be responsible for producing WASH1, the protein that forms an integral part of the endosome-based WASH complex. However, peptide, cDNA and conservation evidence suggests that it is one of the four novel WASH1 genes, *LOC124908094*, that codes for the WASH complex protein. The *LOC124908094* protein, WASH1-20p13, captures almost all the known peptide evidence, is the most similar to conserved WASH1 homologues in other species, and it is just one amino acid different from the protein that would be produced by the GenBank WASH1 cDNA (BC048328.1).

Although the evidence overwhelmingly suggests that WASH1-20p13 is the functional WASH complex protein, it is possible that other WASH1 genes might code for proteins. Five of the 52 tryptic WASH1 peptides recorded PeptideAtlas do not map to WASH1-20p13 and most of these peptides are supported by convincing spectra. However, it is possible that some or all of the non-matching peptides recorded in PeptideAtlas are from common single amino acid variants of *LOC124908094*. In any case, no single WASH1 gene captures all five non-matched peptides and the gene that captures most, *WASH6P*, has a novel N-terminal without peptides and a premature stop codon variant with a major allele frequency of over 0.1%. Full length WASH1 proteins other than WASH1-20p13 have multiple deletions and non-conservative substitutions and analysis using mammalian WASH1 genes found no evidence for positive selection in these human genes [Linardopoulou et al, 2007].

*WASHC1*, the only gene currently annotated as coding in all three of the main human reference gene sets, was thought to be most likely to produce the WASH1 protein because it was the only WASH1 gene that could produce a full length protein. However, even though it is full length, the *WASHC1* protein has 11 single amino acid variants in regions that are highly conserved across tetrapods, and most of these are non-conservative substitutions. In addition to the non-conservative substitutions, there is not a single peptide in PeptideAtlas that uniquely identifies the *WASHC1* protein.

WASH1 genes equivalent to the human *WASH8P* and *LOC124908094* genes are annotated in the subtelomeric regions of chromosomes 12 and chromosome 20 in chimp and bonobo, and chromosome 12 and 20 WASH1 genes have also been found in gorilla [Linardopoulou et al, 2007]. The equivalent human gene on chromosome 12, *WASH8P*, has undergone substantial changes since the divergence from chimpanzee and is likely to be a pseudogene. Phylogenetic analysis, cross species conservation and gene neighbourhoods all confirm that the 11 novel WASH1 paralogues annotated in the human genome appeared in human lineage, and derived from *WASH8P* on chromosome 12 rather than *LOC124908094* on chromosome 20.

The fact that none of the 11 WASH1 paralogues arose from *LOC124908094* strongly suggests that possessing two copies of the functional WASH1 gene is deleterious. Since in this model all 11 novel paralogues came from a gene that was already pseudogenic, it suggests that all 12 WASH1 copies other than *LOC124908094* are pseudogenes, or at least in the process of becoming pseudogenes. An argument against this model in which two copies of a WASH1 coding gene are deleterious is that there are two genes apparently producing full length proteins in both chimpanzee and bonobo.

At present, *WASHC1* on chromosome 9 is annotated as coding and all functional information for the WASH1 protein links to this gene. However, this gene’s history, protein sequence and lack of peptide support suggests that it is probably a pseudogene, even though it does not have an obvious disabling mutation. It will be important to annotate *LOC124908094* as the gene that produces the WASH1 protein in the three main reference databases, not least to make it easier to carry out research into the function of the human WASH complex. Given the importance of the WASH complex in developmental and neurodegenerative disorders [Courtland et al, 2021, Valdmanis et al, 2007], annotating *LOC124908094* as the gene that produces the WASH1 protein will also be essential for mapping clinically relevant variants for these disorders.

Discovering a novel WASH1 coding gene in the regions recently annotated in the T2T genome assembly demonstrates the importance of annotating the whole genome. Although many of the genes added by this new assembly are likely to be pseudogenes, this discovery suggests that there might be other interesting coding genes yet to be discovered in the novel regions annotated by the T2T consortium.

## Acknowledgements

This work was funded by the National Human Genome Research Institute of the National Institutes of Health [grant number U41 HG007234].

## Notes

### Competing Interest Statement

The authors have declared no competing interest.

### Summary of Updates

1. Update author 1 surname. 2. Update first paragraph of introduction 3. Add funding details 4. Add Alekhine et al reference as part introductory paragraph update.

## References

Alekhina, O., Burstein, E., & Billadeau, D. D. (2017). Cellular functions of WASP family proteins at a glance. Journal of cell science, 130, 2235–2241.

Courtland, J. L., Bradshaw, T. W., Waitt, G., Soderblom, E. J., Ho, T., Rajab, A., Vancini, R., Kim, I. H., & Soderling, S. H. (2021). Genetic disruption of WASHC4 drives endo-lysosomal dysfunction and cognitive-movement impairments in mice and humans. eLife, 10, e61590.

Derivery, E., Sousa, C., Gautier, J. J., Lombard, B., Loew, D., & Gautreau, A. (2009). The Arp2/3 activator WASH controls the fission of endosomes through a large multiprotein complex. Developmental cell, 17, 712–723.

Deutsch, E. W., Mendoza, L., Shteynberg, D. D., Hoopmann, M. R., Sun, Z., Eng, J. K., & Moritz, R. L. (2023). Trans-Proteomic Pipeline: Robust Mass Spectrometry-Based Proteomics Data Analysis Suite. Journal of proteome research, 22, 615–624.

Frankish, A., Carbonell-Sala, S., Diekhans, M., Jungreis, I., Loveland, J. E., Mudge, J. M., Sisu, C., Wright, J. C., Arnan, C., Barnes, I., Banerjee, A., Bennett, R., Berry, A., Bignell, A., Boix, C., Calvet, F., Cerdán-Vélez, D., Cunningham, F., Davidson, C., Donaldson, S., … Flicek, P. (2023). GENCODE: reference annotation for the human and mouse genomes in 2023. Nucleic acids research, 51, D942–D949.

Gomez, T. S., & Billadeau, D. D. (2009). A FAM21-containing WASH complex regulates retromer-dependent sorting. Developmental cell, 17, 699–711.

Guindon, S., Dufayard, J.F., Lefort, V., Anisimova, M., Hordijk, W., & Gascuel, O. (2010) New Algorithms and Methods to Estimate Maximum-Likelihood Phylogenies: Assessing the Performance of PhyML 3.0. Systematic Biology, 59, 307–321.

Huang, L., Zhu, P., Xia, P., & Fan, Z. (2016). WASH has a critical role in NK cell cytotoxicity through Lck-mediated phosphorylation. Cell death & disease, 7, e2301.

IJdo, J. W., Baldini, A., Ward, D. C., Reeders, S. T., & Wells, R. A. (1991). Origin of human chromosome 2: an ancestral telomere-telomere fusion. Proceedings of the National Academy of Sciences of the United States of America, 88, 9051–9055.

Kusebauch, U., Deutsch, E. W., Campbell, D. S., Sun, Z., Farrah, T., & Moritz, R. L. (2014). Using PeptideAtlas, SRMAtlas, and PASSEL: Comprehensive Resources for Discovery and Targeted Proteomics. Current protocols in bioinformatics, 46, 13.25.1–13.25.28.

Lefort, V., Longueville, J-E., & Gascuel, O. (2017) SMS: Smart Model Selection in PhyML. Molecular Biology and Evolution, msx149.

Linardopoulou, E. V., Williams, E. M., Fan, Y., Friedman, C., Young, J. M., & Trask, B. J. (2005). Human subtelomeres are hot spots of interchromosomal recombination and segmental duplication. Nature, 437, 94–100.

Linardopoulou, E. V., Parghi, S. S., Friedman, C., Osborn, G. E., Parkhurst, S. M., & Trask, B. J. (2007). Human subtelomeric WASH genes encode a new subclass of the WASP family. PLoS genetics, 3, e237.

Martin, F. J., Amode, M. R., Aneja, A., Austine-Orimoloye, O., Azov, A. G., Barnes, I., Becker, A., Bennett, R., Berry, A., Bhai, J., Bhurji, S. K., Bignell, A., Boddu, S., Branco Lins, P. R., Brooks, L., Ramaraju, S. B., Charkhchi, M., Cockburn, A., Da Rin Fiorretto, L., Davidson, C., … Flicek, P. (2023). Ensembl 2023. Nucleic acids research, 51, D933–D941.

Nurk, S., Koren, S., Rhie, A., Rautiainen, M., Bzikadze, A. V., Mikheenko, A., Vollger, M. R., Altemose, N., Uralsky, L., Gershman, A., Aganezov, S., Hoyt, S. J., Diekhans, M., Logsdon, G. A., Alonge, M., Antonarakis, S. E., Borchers, M., Bouffard, G. G., Brooks, S. Y., Caldas, G. V., … Phillippy, A. M. (2022). The complete sequence of a human genome. Science, 376, 44–53.

Sayers, E. W., Bolton, E. E., Brister, J. R., Canese, K., Chan, J., Comeau, D. C., Farrell, C. M., Feldgarden, M., Fine, A. M., Funk, K., Hatcher, E., Kannan, S., Kelly, C., Kim, S., Klimke, W., Landrum, M. J., Lathrop, S., Lu, Z., Madden, T. L., Malheiro, A., … Sherry, S. T. (2023a). Database resources of the National Center for Biotechnology Information in 2023. Nucleic acids research, 51, D29–D38.

Sayers, E. W., Cavanaugh, M., Clark, K., Pruitt, K. D., Sherry, S. T., Yankie, L., & Karsch-Mizrachi, I. (2023b). GenBank 2023 update. Nucleic acids research, 51, D141–D144.

Strausberg, R. L., Feingold, E. A., Grouse, L. H., Derge, J. G., Klausner, R. D., Collins, F. S., Wagner, L., Shenmen, C. M., Schuler, G. D., Altschul, S. F., Zeeberg, B., Buetow, K. H., Schaefer, C. F., Bhat, N. K., Hopkins, R. F., Jordan, H., Moore, T., Max, S. I., Wang, J., Hsieh, F., … Mammalian Gene Collection Program Team (2002). Generation and initial analysis of more than 15,000 full-length human and mouse cDNA sequences. Proceedings of the National Academy of Sciences of the United States of America, 99, 16899–16903.

UniProt Consortium (2023). UniProt: the Universal Protein Knowledgebase in 2023. Nucleic acids research, 51, D523–D531.

Valdmanis, P. N., Meijer, I. A., Reynolds, A., Lei, A., MacLeod, P., Schlesinger, D., Zatz, M., Reid, E., Dion, P. A., Drapeau, P., & Rouleau, G. A. (2007). Mutations in the KIAA0196 gene at the SPG8 locus cause hereditary spastic paraplegia. American journal of human genetics, 80, 152–161.

Xia, P., Wang, S., Huang, G., Zhu, P., Li, M., Ye, B., Du, Y., & Fan, Z. (2014). WASH is required for the differentiation commitment of hematopoietic stem cells in a c-Myc-dependent manner. The Journal of experimental medicine, 211, 2119–2134.

